# Conservation of mRNA operon formation in the control of the heat shock response in mammalian cells

**DOI:** 10.1101/2024.10.05.616786

**Authors:** Emese Pataki, Jeffrey E. Gerst

## Abstract

Prokaryotic organisms rely on polycistronic transcription (*i.e.* operons) to express multiple mRNAs from a single promoter, enabling rapid and robust responses to stimuli. In yeast, a similar mechanism exists whereby monocistronic mRNAs are selectively assembled into ribonucleoprotein particles, termed RNA operons or transperons, to regulate the expression of genes involved in the same biological process. One example is the heat shock protein (HSP) transperon that confers the eukaryotic heat shock response (HSR), a conserved cellular mechanism that enables organisms to cope with proteotoxic stress. As it was unknown whether transperons exist in higher eukaryotes, we examined whether a similar mechanism operates in mammalian cells. Using both single-molecule fluorescent *in situ* hybridization and RNA pulldown techniques, we show that mammalian HSP mRNAs colocalize and multiplex to form mRNA assemblages during heat stress. These RNP assemblies are dependent on heat shock factor 1 transcriptional regulator and involve both inter- and intrachromosomal interactions among the HSP genes. Bioinformatic analysis identifies a conserved sequence motif within the coding regions of HSP mRNAs and mutational studies in yeast suggest that it is critical for mRNA multiplexing and the HSR. These findings emphasize the evolutionarily conserved nature of heat shock gene regulation across species and suggest that mammalian cells employ RNA operons/transperons in organizing their heat shock response.

## Introduction

Prokaryotic organisms efficiently regulate gene expression through polycistronic transcription units, such as operons, allowing for the simultaneous expression of multiple genes from a single promoter in response to environmental stimuli (Jacob & Monod, 1961). While eukaryotes lack DNA-based operons, it has been suggested they may have evolved analogous mechanisms to coordinate gene expression (*i.e.* RNA operons) (Keene & Tenenbaum, 2002; Mitchell & Parker, 2014). Recent studies have shown that eukaryotic cells utilize various genome organization strategies to regulate gene expression, including chromatin looping, inducible transcriptional condensates, and the formation of “transcriptional factories.” These mechanisms allow for the clustering of genes involved in shared biological pathways, enhancing the efficiency and coordination of gene expression in response to stimuli. For example, in yeast, transcriptional condensates, like the super-enhancers found in mammalian cells, enable the rapid activation of stress-responsive genes (Chowdhary et al., 2019; Kainth et al., 2021).

In yeast, Gross and colleagues demonstrated that heat shock factor 1 (HSF1) mediates inter-chromosomal interactions between heat shock protein (HSP) genes to drive formation of transcriptional condensates with the RNA Pol II and Mediator complexes in yeast (Chowdhary et al., 2022). This interaction results in coordinated gene expression and transcriptional control across different chromosomes. Work by our laboratory extended these findings by identifying the existence of transperons, multiplexed assemblies of co-transcribed mRNAs bound in *trans* that are derived from both intra- and inter-chromosomal interactions, and which encode proteins that function on the same cellular pathway(s) (Nair et al., 2021, 2022). Consistent with the hypothesis of eukaryotic RNA operons, we identified transperons encoding components of the mating pathway of either yeast haplotype, another for mitochondrial outer membrane proteins, and one composed of HSP transcripts (*e.g*. Hsp104, Hsp12, Hsp82, Ssa2, and Ssa4; HSP transperon) that was found to be crucial for the yeast heat shock response (Nair et al., 2021). Notably,transperon assembly and function depends upon histone H4, and its depletion leads to defects in RNA multiplexing and loss of physiological functions (Nair et al., 2021). Transperon formation is reliant on nuclear architecture and histone modifications, which suggests a complex layer of regulation involving chromosomal coalescence (Nair et al., 2022).

From yeast to humans, the same core group of chaperone-encoding genes is induced by heat shock and under the regulation of HSF1 transcription factor, the key regulator of HSR, (Gomez-Pastor et al., 2018; Kurop et al., 2021; Pincus, 2020). Therefore, the HSR is an excellent system to study eukaryotic gene regulation based on its robust induction and conserved components. Despite its name, the HSR is not only sensitive to temperature change, but is also activated by oxidative stress, glucose deprivation and by the presence of misfolded proteins. The HSR relies on a winged helix DNA-binding heat shock transcription factor, HSF1, that recognizes a motif found in the promoter of chaperone genes, (Nelson & Littlefield, 1999; Hentze et al., 2016; Pincus, 2020). Under basal cellular conditions in mammalian cells, inactive HSF1 resides in the cytoplasm forming a complex together with HSP40, HSP70, HSP90 and the TCP1 ring complex (TRiC) (Gomez-Pastor et al., 2018; Kurop et al., 2021). Under stress conditions, HSF1 dissociates from the complex, trimerizes, and translocates to the nucleus where it binds its target genes to initiate transcriptional activation of chaperones including HSP90, HSP70 and small heat shock proteins, which help to refold any denatured proteins and maintain cellular proteostasis. After protein homeostasis has been restored, the excess of HSP70 acts as a negative feedback signal on HSF1, which facilitates its relocation to the cytosol, (Anckar & Sistonen, 2011; Mercier et al., 1999; Webster et al., 2019). Given that the HSR is highly conserved and that HSP genes undergo coalescence upon heat shock, we hypothesized that HSF1-mediated transcription in mammalian cells might also lead to HSP transperon formation to facilitate a coordinated cellular response to heat stress and proteotoxic conditions. By employing single-molecule fluorescent *in situ* hybridization (smFISH), chromatin conformation capture (3C), and single-species RNA pulldown techniques, we report HSP transperon formation in mouse embryo fibroblasts (MEFs) exposed to brief heat shock. Mammalian HSP transperon formation is dependent upon the exposure to heat shock and necessitates the presence of HSF1, as both chromosomal allelic interactions and transperon formation were abolished in cells lacking HSF1. Thus, mRNA operon/transperon formation is conserved in eukaryotes at least with respect to the HSR. Given the complexity of mammalian systems, transperon study may offer significant insights into gene regulation and open up avenues for therapeutic interventions targeting stress response pathways.

## Results

### Inter- and intragenic association between mammalian heat shock genes

Previous work has demonstrated that intergenic associations occur between HSP genes under heat stress leading to transperon formation and the HSR. To investigate whether mammalian HSP genes also exhibit allelic coupling during heat shock, we employed the Chromosome Conformation Capture (3C) technique (Figure 1A) (Lieberman-Aiden et al., 2009). Using DNA samples from mouse embryonic fibroblast cells (MEFs) exposed to heat shock (42°C; 1hr) and PCR, we identified two significant interactions: an intrachromosomal interaction between HSPA1A (HSP70) and HSP90AB1 (HSP90) (Figure 1B-D), and an interchromosomal interaction between HSPA1A and DNAJA1 (HSP40) (Figure 1E-G). In both cases, the N-terminal regions of HSPA1A were involved, suggesting a hotspot for interactions within this region as previously seen in yeast (Nair et al., 2021). Interactions were confirmed by both PCR (Figure 1C, F) and DNA sequencing, and were found to be specific as no interactions were observed between other regions of the genes. Notably, knockout of HSF1 abolished the allelic coupling between HSPA1A and both HSP90AB1 and DNAJA1 (Figure 1D and G, respectively). However, reintroducing HSF1 under a promoter with mild expression restored these interactions (Figure 1D and G). Therefore, we conclude that mouse HSP genes can physically interact and that this allelic coupling is HSF1-dependent. Despite the knowledge that HSP70, HSP90 and HSP40 form a complex together with HSF1, we did not observe intergenic interaction between HSP90AB1 and DNAJA1 (Supplementary Figure 1). This might be due to the existence of multiple HSP isoforms within each HSP family in mammalian cells. Transcriptome sequencing of MEFs in our lab has revealed that >30 different HSP isoforms are transcribed under non-heat shock conditions alone, with varying degrees of expression (Dasgupta et al., 2023; Source Data 1 file).

**Figure 1.**
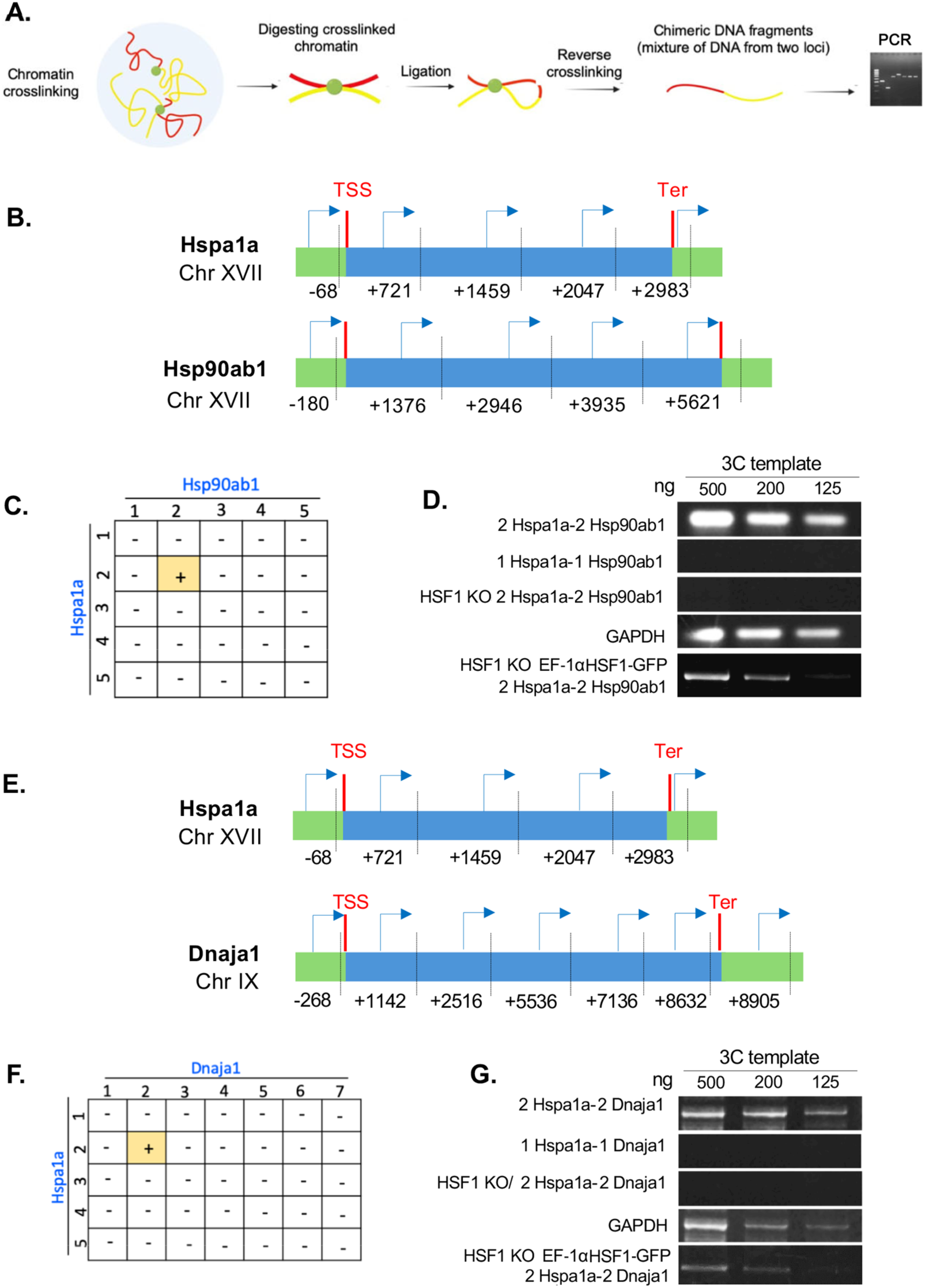
Intergenic interactions of mammalian HSP genes. (A) Visual outline of the Chromosome conformation capture (3C) technique. (B, E) Graphic illustration of the HSPA1A and HSP90AB1 or HSPA1A and DNAJA1 genes, and the oligonucleotides used for their amplification from 3C DNA samples. Coordinates correspond to TaqI sites (shown as vertical dashed bars); site numbering is relative to ATG (+1). Forward primers used for 3C analysis were sense-strand identical (arrows) and positioned proximal to TaqI sites as indicated. Primers are numbered to distinguish the pairs used in the PCR reactions shown in (D, G). 5’UTRs, ORFs, and 3’UTRs are color-coded, as indicated. Transcription start sites (TSS) and termination sites (Ter) are indicated. (C, F) Matrices summarizing the intergenic associations of represented genes as determined by 3C-PCR. Primer pairs corresponding to the different genes listed in (B, E) were used in PCR reactions. ‘+’ indicates PCR amplification and the interaction between genes. ‘–’ indicates no amplification. (D, G) PCR products derived reactions using the indicated primer pairs (to the genes shown in B, E) and 3C-processed DNA were electrophoresed on agarose gels (1%) and visualized by ethidium bromide staining. Lanes represent the 3C-PCR output using the indicated concentrations of DNA template (ng DNA). *GAPDH* primers were used as a control for the PCR reaction.

### HSP mRNAs co-localize in MEF cells upon exposure to heat shock

Since HSPA1A and HSP90AB1 physically interact on the same chromosome and undergo allelic coupling, we investigated whether these genes also interact at the RNA level. To address this, we performed smFISH using specific fluorescent oligonucleotide probes (Figure 2; Supplementary File 1). First, to assess HSP mRNA expression levels in MEFs, we extracted RNA from cells maintained at 37°C or exposed to heat shock (42°C; 1 hr). As shown in Supplementary Figure 2, several heat shock mRNAs were expressed under normal growth conditions, but their expression levels strongly increased following heat stress. In contrast, HSP mRNA levels were significantly reduced after heat shock in HSF1^-/-^ knockout cells, with minimal expression observed under normal growth conditions (Supplementary Figure 2). We used smFISH to examine for mRNA interactions, targeting HSPA1A mRNA with sequence-specific probes labeled with Quasar® 670 and HSP90AB1 mRNA with probes labeled with Quasar® 570 (Figure 2A). The number of mRNA signals was detected and quantified using FISH-quant (Figure 2B) (Mueller et al., 2013). No HSPA1A mRNA signals were observed in cells not exposed to heat shock, while HSP90AB1 mRNA spots were observed (not shown). In contrast, both HSPA1A mRNA and HSP90AB1 mRNA signals were detected upon heat shock (Figure 2A). Quantification revealed that 17% of HSPA1A and HSP90AB1 mRNAs were co-localized (Figure 2B), the partial overlap perhaps explained by the constitutive expression of HSP90AB1 and suggesting that co-localization occurs only after the induction of HSPA1A. As a control, we examined RNA co-localization between β-actin mRNA and either HSPA1A or HSP90AB1 but did not observe any interaction. As both HSPA1A and HSP90AB1 reside on Chr. XVII, we examined whether HSPA1A mRNA could co-localize with another HSP mRNA when the genes are located on two different chromosomes (*e.g.* HSPA1a on Chr. XVII and HSPA8 on Chr. IX). Representative smFISH images (Figure 2C) show the labeled HSPA1a and HSPA8 mRNAs and their level of co-localization (Figure 2D). The results show that HSPA1A mRNAs overlap with HSPA8 mRNAs with 26% co-localization upon heat shock. Again, little interaction was observed between either HSPA1a or HSPA8 mRNA and β-actin mRNA. Thus, the HSP mRNA interactions observed appear specific.

**Figure 2.**
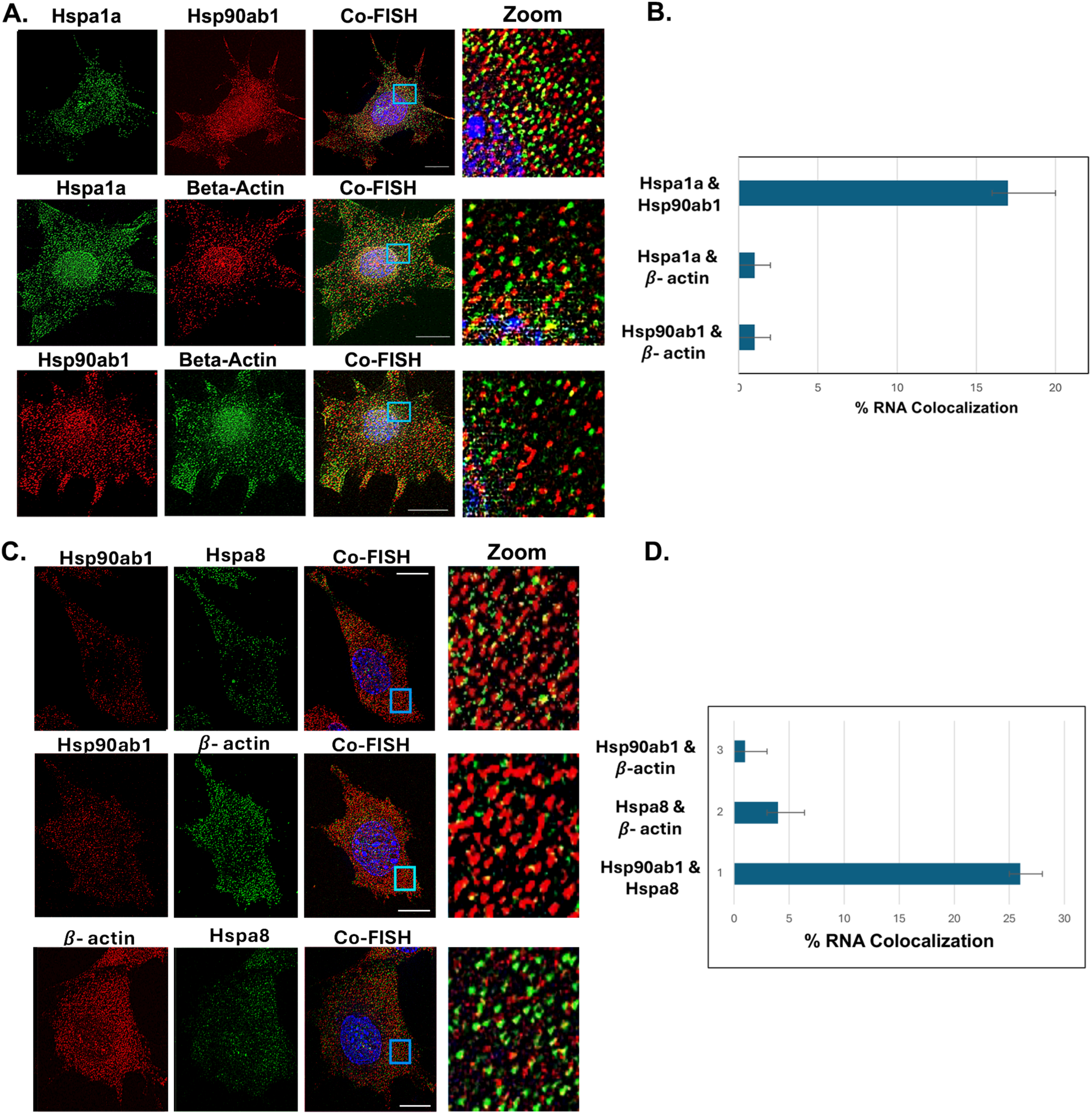
HSP mRNAs co-localize upon heat shock. (A, C) Representative single-molecule fluorescence *in situ* hybridization (smFISH) images of MEFs that underwent heat shock (HS) or did not (NHS). The cells were processed for smFISH labeling using sequence-specific FISH probes complementary to HSPA1A, HSP90AB1 and HSPA8 mRNAs, prior to labeling with DAPI (blue). HSPA1A and HSPA8 mRNAs were labeled with Quasar® 670-labeled oligos (green channel), while HSP90AB1 mRNA was labeled with Quasar® 570-labeled oligos (red channel). Images were filtered and analyzed using Fish-Quant. *Co-FISH* indicates the merge of all three signals. *Zoom* indicates a close-up from the merged signals. (B, D) Histogram of the data obtained from three biological replicates.

### mRNAs encoding mammalian HSPs multiplex to form a heat shock-induced transperon

To investigate whether mRNAs encoding mammalian HSPs multiplex to form transperons we employed an RNA-specific pulldown assay created by Torres *et al*. (2018) (Torres et al., 2018) with some modifications, as detailed in the *Materials and Methods*. We designed biotin-labeled DNA oligonucleotides complementary to HSPA1A and hybridized them with whole cell extracts derived from formaldehyde-cross-linked WT and HSF1^-/-^ knockout (KO) cells that were incubated at either 42°C to induce heat shock or maintained at 37°C as a control. In addition, we further assessed the role of HSF1 by comparing multiplexing in WT and HSF1^-/-^ KO cells treated under these same conditions. Next, we used immobilized streptavidin to precipitate the HSPA1A RNA followed by reverse transcription-PCR (RT-PCR) to detect co-precipitated mRNAs after normalization for expression (Figure 3A).

**Figure 3.**
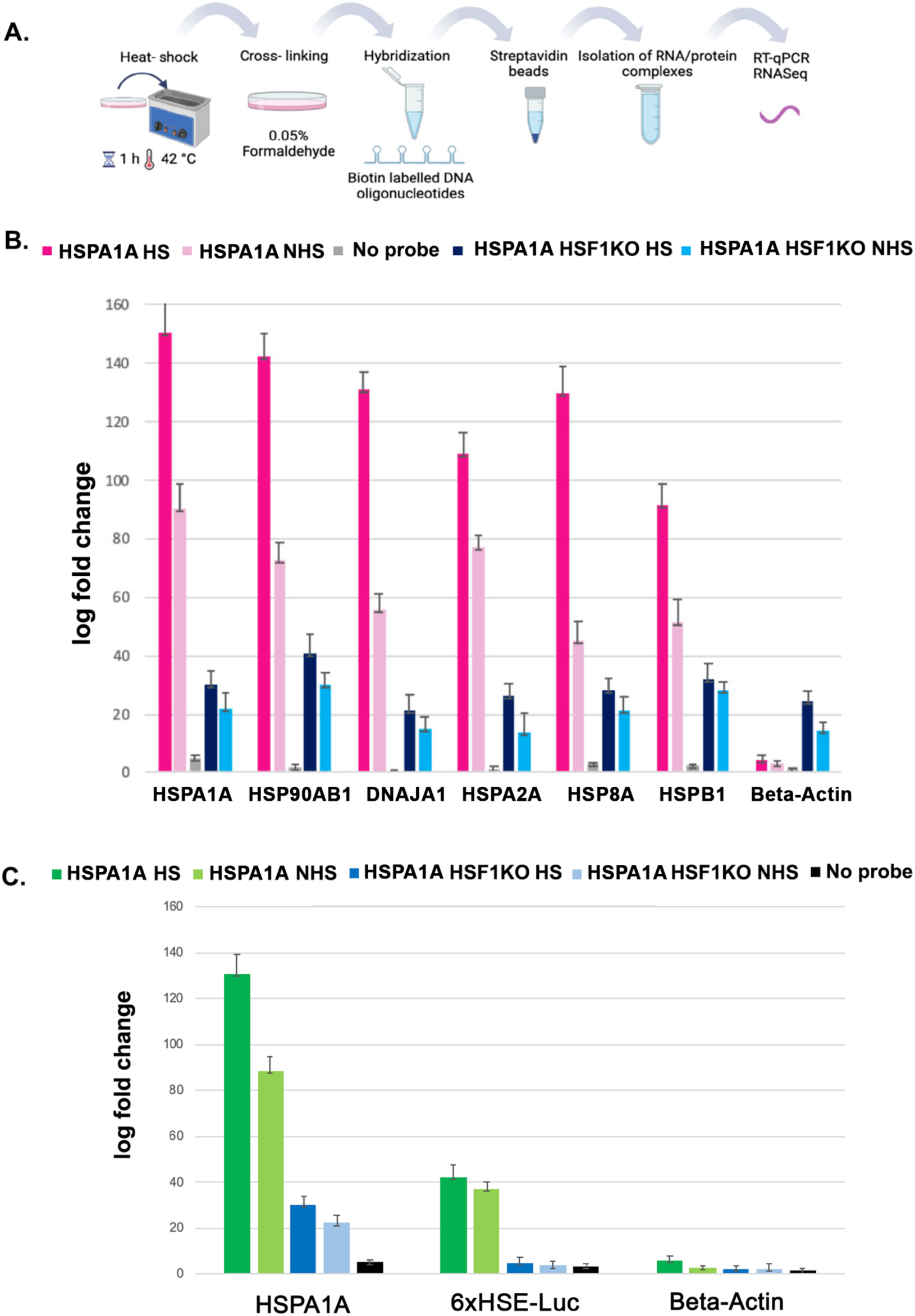
HSF1 and the HSP promoter containing a HSRE drive HSP RNA multiplexing. (A) Schematic of the RNA pulldown technique using biotin-labeled DNA oligonucleotides. (B) WT MEF and HSF1^-/-^ KO MEFs were either exposed to heat shock (1h; 42℃; HS) or maintained at 37℃ (NHS) prior to fixation and the RNA pulldown procedure. RT-PCR was performed using specific oligos against the listed genes and quantified after normalization for expression. Three biological replicates were performed. The *No probe* condition indicates that no biotin-labeled DNA oligos were added to the cells, as a negative control. (C) 6xHSE-RLuc plasmid was transiently expressed in WT and HSF1^-/-^ KO MEF cells and either subjected to heat shock at 42℃ for 1 hr or maintained at 37 ℃ prior to fixation and pulldown procedure. RT-PCR was performed using specific oligos against the listed genes and quantified after normalization for expression Three biological replicates were performed.

Notably, the mRNAs of HSP genes previously shown to undergo intergenic interactions and mRNA co-localization, such as HSP90AB1, DNAJA1, and HSPA8 (Figures 1 and 2), along with HSPB1, showed co-precipitation with HSPA1A mRNA that was more robust upon the exposure of cells to heat shock (Figure 3B). Furthermore, HSF1 was once again identified as a critical regulator of RNA multiplexing, as in its absence the amount of HSP mRNAs that co-precipitated with HSPA1A RNA was greatly reduced. These results demonstrate that the HSP RNA transperon is conserved in evolution.

Since HSF1 plays an essential role in promoting the formation of heat shock transperon and the HSR in both yeast and mammalian cells, we hypothesized that presence of a heat shock responsive element (HSRE) in any given RNA might be sufficient to confer allelic coalescence, co-transcription, and assembly into the RNA operon upon heat shock. This is because active oligomeric HSF1 has been suggested to undergo liquid phase-phase separation on the promoters of HSP genes to form a condensate that allows for efficient PolII transcription (Chowdhary et al., 2022) To test the possibility that a HSRE is necessary and sufficient to confer RNA incorporation into the HSP transperon, we transfected MEFs with plasmids either expressing pGLuc or 6xHSE-Rluc, the latter containing six copies of the HSRE, into both WT and HSF1^- /-^ KO cells and precipitated HSPA1A RNA. Upon RT-PCR analysis, we found that HSRE- luciferase RNA was incorporated into the RNA multiplex in a heat shock- and HSF1-dependent manner. In contrast, β-actin mRNA was not incorporated under any condition. As expected, the GLuc luciferase gene alone lacking the HSREs did not appear in the multiplex (Supplementary Figure 3.). Thus, the HSREs appear to confer mRNA participation in multiplexing, indicating an essential role for this transcription factor in the control of transperon formation.

### A conserved sequence motif within the coding region of heat shock mRNAs promotes RNA multiplexing

Since the HSR genes are conserved across species from yeast to humans, we performed a bioinformatic analysis to identify recognizable motifs in their sequences (Bailey et al., 2009). Using MEME-ChIP analysis (https://meme-suite.org/meme/tools/meme-chip), we identified a conserved motif within the coding regions of *S. cerevisiae* and *Mus musculus* HSP mRNAs, represented as a sequence logo based on nucleotide composition (Figure 4A). To determine whether this motif contributes to RNA multiplexing, we generated a mutated version of the yeast *HSP82* gene by substituting four conserved nucleotides within the motif with synonymous mutations that did not alter the amino acid sequence (*hsp82^mut^*; Figure 4B). Structural analysis using RNA-fold revealed that *hsp82^mut^* was distorted compared to the native *HSP82* sequence (Figure 4C). In addition, we examined the phenotype of the *hsp82^mut^* strain under temperature and osmotic stress conditions and found that it closely resembled that of an *hsp82Δ* deletion mutant (Figure 4D), indicating that these mutations resulted in a loss of function.

**Figure 4.**
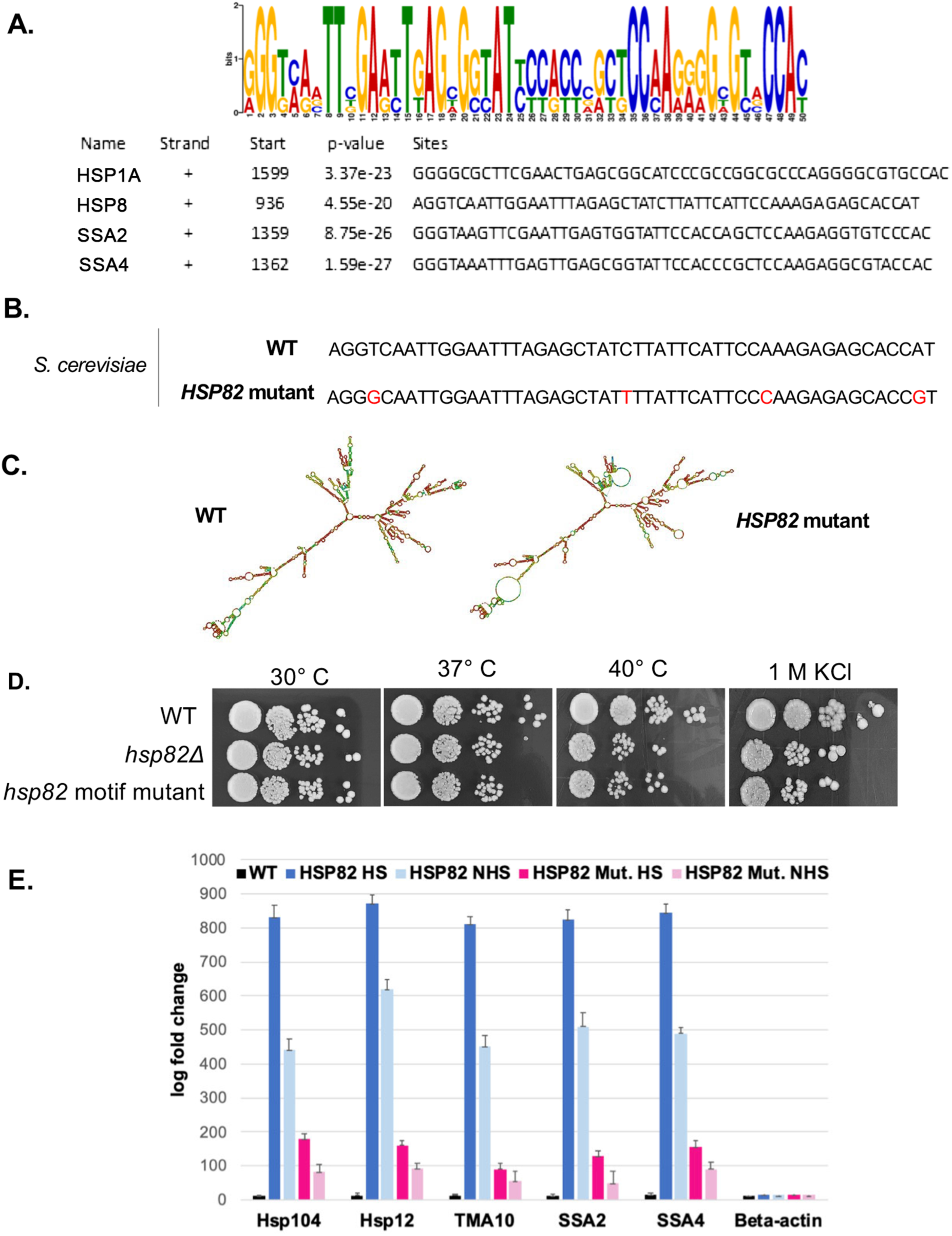
A conserved motif in the HSP mRNA coding region facilitates HSP transperon assembly and function in yeast. (A) A conserved sequence motif in the coding region is common to the *Mus musculus HSPA1A* and *HSP8* genes and the *S. cerevisiae SSA2* and *SSA4* HSP genes. MEME-ChIP analysis of the sequences of all four heat shock protein mRNAs was performed and revealed a consensus motif in the coding regions, shown schematically as a sequence logo based on nucleotide representation. (B) Shown are the *S. cerevisiae HSP82* (WT) and mutant forms in which the motif was altered within the coding region to change conserved nucleotides without altering the amino acid sequence. (C) The *HSP82* mutant has an altered secondary structure. RNA-fold was used to model the native and mutant versions of the *HSP82* mRNA. (D) Mutation of the consensus motif in *HSP82* leads to decreased growth under high temperature and osmotic stress similarly to the *hsp82Δ* mutant. Serial dilutions of cells bearing WT *HSP82* or the *HSP82* motif mutant, as well as *hsp82Δ* cells were spotted onto YPD medium with or without 1M KCl and incubated for 2 days at the indicated temperatures. (E) Mutation of the consensus motif in HSP82 leads to a decrease in HSP mRNA multiplexing. Cells expressing MS2 aptamer-tagged *HSP82* or *HSP82* motif mutant (*HSP82 mut*) were either exposed to heat shock (10 min; 40℃; HS) or maintained at 30℃ (NHS) prior to fixation and RaPID RNA pulldowns and qRT-PCR. Three biological replicates were performed.

Finally, we used RaPID, an RNA pulldown technique from our lab that uses the MS2 coat protein fused to GFP and streptavidin-binding peptide to precipitate MS2 aptamer-tagged mRNAs (Slobodin & Gerst, 2011; Zabezhinsky et al., 2016; Nair et al., 2021) from cells bearing the *HSP82* and *hsp82^mut^* genes tagged at the *HSP82* locus with twelve MS2 aptamer sequences. Upon RNA pulldown using immobilized streptavidin, we found that HSP mRNAs previously shown to co-precipitate with WT *HSP82* mRNA (*e.g. HSP104, HSP12, TMA10, SSA2,* and *SSA4* mRNAs) (Nair et al., 2021) were significantly reduced in *hsp82^mut^* cells as compared to WT cells (Figure 4E). These findings suggest that the conserved motif plays a crucial role in heat shock-induced HSP mRNA multiplexing and their ability to confer the HSR (Figure 4D).

## Discussion

The present study explores the regulation of mammalian HSP gene co-transcription and mRNA multiplexing during the HSR, extending our earlier findings from yeast now to mammalian systems. The proposed mechanism is summarized in Figure 5, which visually illustrates the interactions and processes identified in this study. By demonstrating that HSP genes undergo HSF1-dependent interactions upon heat shock, subsequent mRNA assembly into RNA operons, and mRNA colocalization in the cytosol, our results highlight a complex evolutionarily conserved regulatory mechanism that governs stress-responsive gene expression.

**Figure 5.**
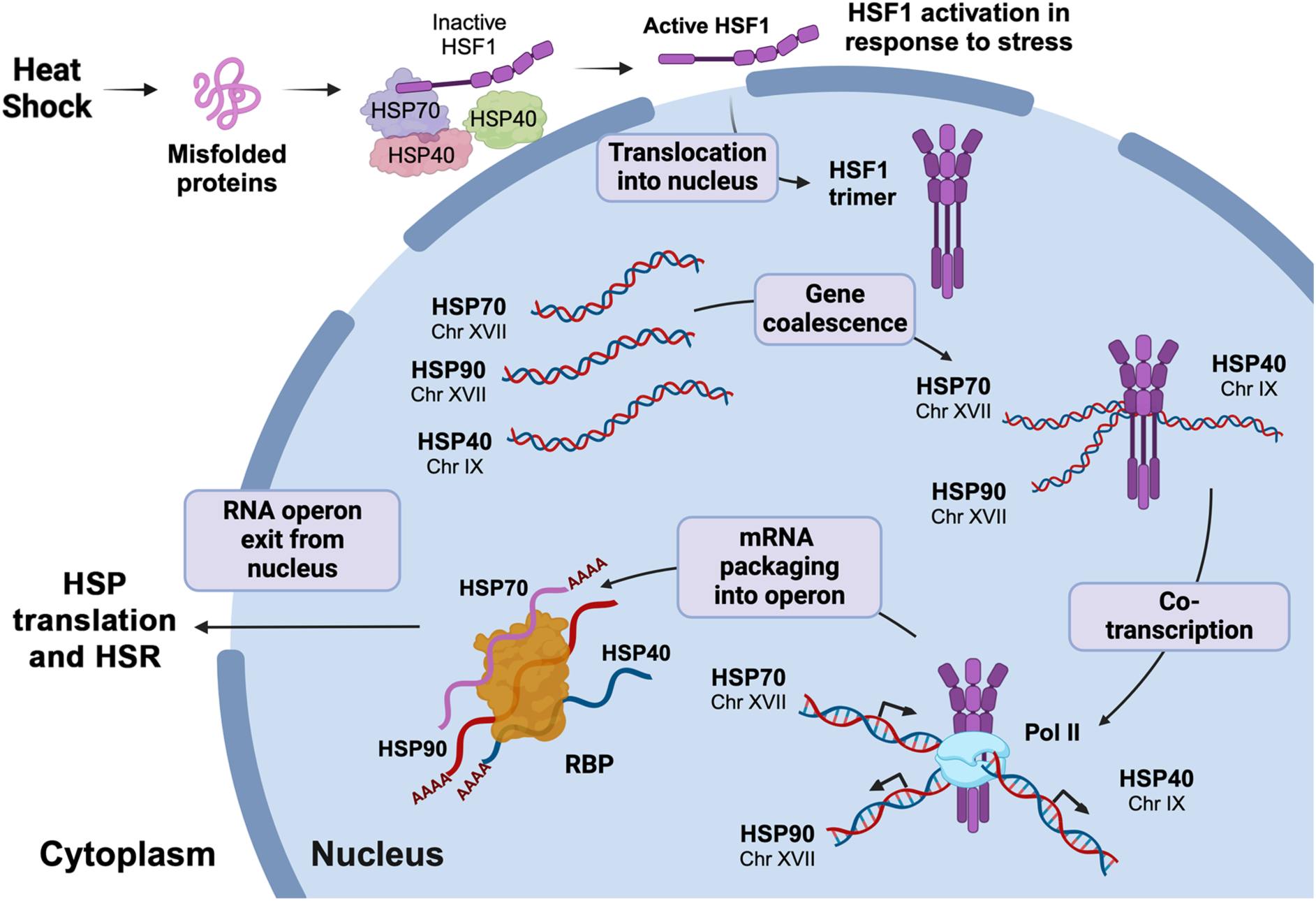
HSP mRNA multiplexing and operon formation is driven by HSF1-mediated HSP gene coalescence and co-transcription. In mammalian cells, inactive HSF1 monomers are retained in the cytoplasm in a complex with regulatory chaperones, such as HSP70, HSP90, and HSP40. In response to stress (*e.g.* heat shock), HSF1 dissociates from this inhibitory complex, trimerizes, and translocates to the nucleus. HSF1 then binds to heat shock elements (HSEs) of target HSP genes and via liquid-liquid phase separation mediates gene coalescence. HSF1-mediated gene coupling allows for interactions with RNA Pol II and the Mediator complex (latter not shown) and leads to coordinated transcription of the HSP genes and subsequent assembly HSP mRNAs into a ribonucleoprotein complex, the heat shock response RNA operon (or transperon). Involvement of yet unknown RNA-binding proteins (RBP) is likely to play a central role in operon assembly. The RNA operon exits the nucleus and upon translation leads to the heat shock response. Upon translation of the HSPs and restoration of proteostasis, HSF1 is transported back to the cytoplasm and returned to its inactive state (not shown).

We first confirmed that mammalian HSP genes engage in physical interactions, both between alleles on the same chromosome and across different chromosomes, during heat shock. Using 3C analysis, we identified significant interactions between HSPA1A (HSP70) and HSP90AB1 (HSP90), as well as between HSPA1A and DNAJA1 (HSP40), both of which were disrupted upon HSF1 gene knockout (Figure 1). The HSF1-dependent coupling of these genes is reminiscent of transcriptional condensates observed in yeast, (Chowdhary et al., 2022) suggesting that mammalian cells utilize a similar mechanism to coordinate HSP gene expression across different genomic loci during stress, and as previously shown by (Zhang et al., 2022). Consequently, we found that HSP mRNAs exhibit colocalization upon heat shock, a feature also mediated by HSF1 (Figure 2). Using smFISH, we detected a substantial increase in colocalization between HSPA1A and HSP90AB1 mRNAs, as well as between the HSPA1A and HSPA8 mRNAs under these conditions. Our findings support the hypothesis that akin to yeast transperons, mammalian cells also form heat-induced RNA-protein complexes (RNPs) that regulate the spatial organization of mRNA molecules involved in stress responses. This idea is further supported by RNA pulldown assays which demonstrated that mammalian HSP mRNAs multiplex to form an HSF1-dependent RNP particle under heat shock conditions. Specifically, HSPA1A mRNA co-precipitated other HSP mRNAs, such as HSP90AB1, DNAJA1, and HSPA8, with this association significantly enhanced after heat shock. The absence of HSF1 drastically reduced the formation of these complexes, underscoring the central role of HSF1 in orchestrating mRNA multiplexing during stress. These results align with our previous work in yeast, where HSF1 was shown to drive intergenic associations and mRNA multiplexing during the HSR (Nair et al., 2021).

In addition to demonstrating the conservation of this process, we also identified a conserved sequence motif within the coding region of heat shock mRNAs that appears to facilitate the assembly of these RNPs (Figure 4). Through mutational analysis of *HSP82* in yeast, we showed that disruption of this motif not only altered RNA structure but also impaired the formation of multiplexed mRNAs, leading to a stress response phenotype that closely resembled that of an *hsp821′* deletion mutant. These findings suggest that the conserved motif plays a pivotal role in mediating the formation of the stress-responsive HSP mRNA operon and its function.

Taken together, our data suggest that mammalian cells employ a highly coordinated mechanism to regulate heat shock gene expression through HSF1-mediated gene interactions, mRNA co-transcription, and the formation of RNP complexes (Figure 5). This work expands our understanding of eukaryotic gene regulation under stress conditions, offering a model that draws parallels between prokaryotic operons and eukaryotic transperons. Moreover, the identification of a conserved motif that facilitates mRNA multiplexing provides a new layer of insight into how eukaryotic cells ensure efficient, synchronized responses to environmental stress. Demonstration of transperon formation in yeast and now in higher eukaryotes is an important step in understanding combinatorial gene expression and its role in coordinating cellular processes. Given the greater complexity of mammalian cells and growing need for new medical intervention strategies, this work may shed new light on basic aspects of gene regulation and be critical for our understanding and ability to manipulate the HSR pathway, which serves a critical function to relieve stress and refold denatured proteins in both normal and stressed cells. Specifically, the HSPs within the HSR are responsible for the folding, activation, and assembly of target proteins as ubiquitous molecular chaperones. HSP overexpression and inhibition can impact the regulation of apoptosis and therefore is targeted by a wide variety of therapies against pathological conditions including cancer, cellular aging, and senescence. It has not escaped our knowledge that interventions that reduce HSP transperon formation might also constitute an effective therapeutic strategy.

## Materials and Methods

### Mammalian Cell Experiments

#### Cell lines and plasmids used in this study

Plasmids used in this study are listed in Supplementary File 1. Immortalized HSF^-/-^ MEFs (Levi-Galibov et al., 2020) were a gift from Ruth Scherz-Shouval (Weizmann Institute of Science, Israel).

Human HSF1-GFP lentivirus vector was a gift from Maria Vera Ugalde (McGill University, Canada). To create EF1α-HSF1-GFP (Addgene plasmid #227820), the UbC promoter driving the expression was replaced by the stronger EF1α promoter, which was PCR-amplified from pTwist-EF1α/Puro plasmid (Twist Bioscience), and subcloned using restriction enzymes *Afe*I and *Not*I.

EF1α-HSF1-GFP lentivirus particles were produced by transiently transfecting the expression plasmid with packaging plasmids, VSVG, RRE and Rev (Addgene plasmids #12259, 12251, and 12253, respectively) into HEK293T cells using calcium phosphate, and allowing the cells to grow for 72 hrs. The virus-containing medium was harvested and concentrated with the Lenti-X concentrator (Clontech), per the manufacturer’s instructions. Viral particles were resuspended in complete DMEM, aliquoted and stored at −80°C until infection.

For stable cell line generation, HSF1^-/-^ MEFs were seeded in 12-well plates and were exposed to the viral particles in serum-free media containing 6 µg/ml polybrene (Sigma) for 2 hours with occasional shaking followed by addition of complete media. Cells with high expression of the fluorescent reporter were selected by FACS sorting (BD Biosciences Aria III).

#### Transient transfection

Transient expression of the luciferase gene and the conserved heat shock response element was carried out using the jetPRIME Transfection Reagent kit. The pGLuc and 6xHSE-Rluc plasmids (gifts from Thomas Czerny) (Ortner et al., 2015) carrying 6 repeats of the conserved heat shock element were transiently co-transfected into WT MEFs or into HSF1^-/-^ mutant MEFs, before subjecting the cells to heat stress and the RNA pulldown procedure.

#### Chromosome conformation capture (3C)

The quantitative chromosome conformation capture method, TaqI-3C, (Chowdhary et al., 2019; Hagège et al., 2007) was employed. MEF cells were first cultured at 37° C to 10^7^ cells in 10 cm plates and either maintained at that temperature or heat shocked for 1hr at 42°C before being crosslinked with 1% formaldehyde for 10 min. Then 125mM glycine was added at room temperature for 5 min to stop the crosslinking. The cells were centrifuged at 1700 *x g* for 1 min and then subjected to lysis in FA lysis buffer (50 mM HEPES at pH 7.9, 140 mM NaCl, 1% Triton X-100, 0.1% sodium deoxycholate, 1 mM EDTA, 1 mM PMSF) on ice for 20 min. Cell lysates were collected and centrifuged for 1 min at 1700 x *g.* The supernatant was transferred to new Eppendorf tube and centrifuged for 10 min at 11,000 x *g* at 4℃. After centrifugation, a thin translucent layer of chromatin was observed on the top of the pellet of cell debris. The supernatant was discarded, and the pellet resuspended in 1 ml FA lysis buffer. The resuspended material was centrifuged at 13,000 x *g* for 10 min at 4°C and the resulting pellet resuspended in 500 μl of 1.2x TaqI restriction enzyme buffer. Next, 7.5 μl of 20% (w/v) SDS (final concentration of 0.3% SDS) was added and the sample incubated for 1hr at 37°C with shaking (900 rpm). Then, 50 μl of 20% (v/v) Triton X-100 (final concentration = 1.8%) was added and the sample further incubated for 1 h at 37°C with shaking (900 rpm). An undigested sample of genomic DNA (50 μl aliquot) was removed and stored at −20°C to determine enzyme digestion efficiency. To the remaining sample, 200 U of TaqI (New England Biolabs) was added and incubated at 60°C for overnight. The next day, 150 μl of digested sample was heat-inactivated at 80°C for 20 min in the presence of added SDS (24 μl 10% SDS; final concentration = 1.7%). To samples of digested material (174 µl each), 626 µl of 1.15x ligation buffer and 80 µl of 10% Triton X-100 (final concentration of 1%) were added and incubated at 37°C for 1 h, while shaking gently. Proximity ligation in the sample was performed using 100 U of added T4 DNA ligase (New England Biolabs) at 16°C for 16 hr. The ligated samples were then digested with RNase (final concentration of 11 ng/μl; RNaseA and 28 U/μl RNAseT1; ThermoScientific) at 37°C for 20 min. Proteinase K (final concentration of 56 ng/μl; Sigma Aldrich) digestion was performed at 65°C for 1 h (note: final concentration of SDS = 0.4%). The 3C DNA template was extracted twice using an equal volume of phenol-chloroform (1:1 TE-saturated phenol-chloroform v/v), followed by extraction with an equal volume of phenol-chloroform-isoamyl alcohol (25:24:1 v/v), followed by extraction with an equal volume of chloroform-isoamyl alcohol (24:1 v/v). The DNA was precipitated in the presence of 2 µl of glycogen (20 mg/ml), sodium acetate (0.3 M final concentration, pH 5.2), and 2.5 volumes of ethanol at RT overnight. The 3C DNA template obtained was stored at −20°C. DNA concentration was determined by absorption spectroscopy at 260 nm. Typically, 125–500 ng of the 3C DNA template was used in the PCR reactions. PCR products were eluted from agarose gels and sequenced for confirmation.

#### Single molecule fluorescent in situ hybridization (smFISH)

smFISH in mammalian cells was carried out according to a previously described protocol (Haimovich et al., 2017; Haimovich & Gerst, 2018, 2019). A set of Quasar (Q) 670-labeled FISH probes to detect HSPA1A or HSPA8 mRNA and Quasar (Q) 570-labeled FISH probes to HSP90AB1 mRNA were obtained from Stellaris. The sequences of the labeled nucleotides is provided in Supplementary File 1. MEF cells were either maintained at 37 ℃ or subjected to heat shock for 1 h at 42℃ followed by 1 h incubation at 37℃ to allow the cells to recover.

#### Widefield imaging

Images of smFISH experiments were captured using a Zeiss AxioObserver Z1 DuoLink dual camera imaging system equipped with an Illuminator HXP 120 V light source and Plan Apochromat 100×1.4 NA oil-immersion objective. Thirty 0.2 μm step *z*-stack images were taken for smFISH. For each individual experiment between 100-150 cells were scored for mRNA and analyzed using the Fish-Quant program (Mueller et al., 2013) (https://bitbucket.org/muellerflorian/fish_quant). Co-localization between the two channels was calculated as a linear assignment problem solved with the Hungarian algorithm. We used Matlab functions hungarianlinker^2^ and munkres^3^ developed by Mueller et al. (2013) for this purpose. We also crosschecked co-localization (defined by overlap between the signals) manually for verification. For statistical analyses, at least three replicate experiments were carried out.

#### RNA pulldown using biotin-labeled DNA oligonucleotides

RNA pulldown using biotin-labeled oligonucleotides was performed following the protocol described by Torres *et al*. (2018). Nine DNA oligonucleotide probes of 25 bases which display strong affinity for the RNA of interest (*e.g.* HSPA1A) were designed as described and obtained as 3’-biotinylated oligos from Sigma. Next, 1×10^7^ cells maintained either at 37°C or shifted for 1hr to 42°C were fixed using freshly prepared 0.1 % paraformaldehyde solution in PBS (10 ml/ culture plate) under agitation for 10 min at room temperature. Paraformaldehyde was quenched by adding 1/10 volume of glycine 1.25M and incubated for 5 min at room temperature. Cells were rinsed two times (5 min each) with PBS. Cells were collected and centrifuged at 510 g at 4 ℃ for 5 min. The pellet was lysed for 15 min on ice in Lysis Buffer (50mM Tris-HCl ph. 7,0, 10mM EDTA, 1% SDS supplemented with 200U/mL of a RNase inhibitor solution, and a cocktail of protease inhibitors at 5µl/ml) (cOmplete, EDTA-free protease inhibitor cocktail tablets, Sigma). Cells were centrifuged for 5 min at 12,000 x *g* at 4℃. Supernatants were hybridized with 100pmol of each of the nine biotinylated oligonucleotides in 2 volumes of hybridization buffer (50mM Tris-HCl ph 7,0, 750 mM NaCl, 1 mM EDTA, 1% SDS, 10 % formamide. Prior to adding the probes, 20 µl from each sample were saved as input samples. Hybridization was allowed to proceed for 4-6 hrs at 30℃. Following hybridization, 50 µl of streptavidin-Sepharose high performance beads (Sigma), supplemented with 200 U/mL of RNAase inhibitor solution and a cocktail of protease inhibitors (cOmplete, EDTA-free protease inhibitor cocktail tablets, Sigma) (5 µl/ml) were added to the samples and were incubated overnight under moderate agitation at room temperature. Beads were washed 5 times for 5 min agitation at room temperature with 900 µl of wash buffer (SDS 0.5%, SSC 2x). After the last wash, 95 µl of Proteinase K buffer (10mM Tris-HCl pH 7.0, 100 mM NaCl, 1 mM EDTA, 0.5% SDS) supplemented with 5 µl of proteinase-K (20mg/mL) was added to the pulldown samples and 75 µl of supplemented Proteinase-K buffer to the previously saved input samples. Samples were incubated at 50 ℃ for 45 min then at 95℃ for 10 min. Samples were placed on ice for 3 min, then total RNA was extracted using Nucleospin RNA Mini kit (Macherey Nagel) and RNA integrity checked using Agilent Tapestation 2100, while the RNA concentration was measured using NanoDrop microvolume spectrophotometer and Qubit 2.0 BR assay. Reverse transcription using XY RT kit followed by qPCR using specific primers (Supplementary file).

### Yeast Cell Experiments

#### Genetic manipulations and plasmids

Yeast strains created for this study were derived from WT BY4741 cells (see Supplementary File 1 for strain list and genotype). Standard LiOAc-based protocols were employed for the introduction of plasmids and PCR products into yeast. For the RaPID RNA pulldown experiment (Slobodin & Gerst, 2011), cells were grown at 30℃ to mid-log phase (O.D._600_=∼1) in a standard rich growth medium containing 2% glucose (YPD) or in a synthetic medium containing 2% glucose (*e.g*. synthetic complete [SC]) and selective drop-out medium lacking amino acid. Induction of the methionine starvation-inducible plasmid (pUG36-MS2-CP-GFP(x3) was performed by transferring cells to mid-log phase to a synthetic medium lacking methionine and subsequent growth of the cells for 1 hr with shaking at 30℃. Any genomic manipulation used for creating the strains for the RaPID pulldown experiment were done following the protocol detailed in (Haim-Vilmovsky & Gerst, 2009). Cells were grown at 30℃ or heat shocked at 50℃ for 10 min- see below for the RaPID protocol employed.

For growth tests on plates, 2x 10^7^ yeast cells were grown to mid-log phase, normalized for absorbance at O.D._600_, serially diluted 1:10, plated by drops onto solid medium, and grown at the indicated temperatures for 48hrs before photodocumentation. Three biological replicas were performed for each growth test.

Chromosomal integration of the four point mutations into the conserved HSP motif sequence of *HSP82* was executed using standard techniques, based on homologous recombination. We first obtained a fragment containing the four point mutations using wild-type genomic DNA as template for two separate PCR reactions. One reaction used primer #1 (5’ CCTCTCTCAACACAGTAATCCATAAAC) together with primer #3 (5’CTCCGTTGAAGG**<u>G</u>**CAATTGGAATTTAGAGCTAT**<u>T</u>**TTATTCATTCC**<u>C</u>**AAGAGAGC ACC**<u>G</u>**TTCGACTTGT) and a second reaction used primer #2 (5’ ATGAGGATGAAGAAACAGAGACTG) together with primer #4 (5’ ACAAGTCGAA**<u>C</u>**GGTGCTCTCTT**<u>G</u>**GGAATGAATAA**<u>A</u>**ATAGCTCTAAATTCCAATTG**<u>C</u>**CCTTCAACGGAG). The underlined nucleotide bases indicate the site we inserted the silent point mutations. For the PCR reaction, 34 cycles of PCR were performed with the following temperature profile: 94°C 1 min, 54°C 1 min, 72°C 2 min. The resulting two PCR fragments were stitched together using 1μl of PCR product from each PCR reaction as template and primers #251 and #252. PCR amplification was achieved by 34 cycles of PCR with the following temperature profile: 94°C 1 min, 48°C 1 min, 72°C 2 min. The PCR product was transformed into the *HSP82::URA3* strain and plated onto 5-FOA plates for selection against *URA3* cells to yield *hsp82^mut^::ura3* cells. Genomic DNA was isolated from cells that grew in the presence of 5-FOA and were further subjected to DNA sequence analysis to ensure the presence of the point mutations at the correct locus. The *HSP82::URA3* strain was created by amplifying the *URA3* gene from pCg plasmid by PCR using primers: Forward 5’ GTCCTATAAACAAAAGCACAAACAAACACGCAAAGATATG*CACAGGAAACAGCTAT GACC* and Reverse 5’ TTTTGTTTATAACCTATTCAAGGCCATGATGTTCTACCTA*GTTGTAAAACGACGGCCAGT*. The PCR product was transformed into BY4741 cells and plated onto minimal media lacking uracil. Integration of the *URA3* gene was confirmed by PCR with primers: Forward 5’ ATAACTTAGCTTGCGTGTTGCGT and Reverse 5’GTACGAACATCCAATGAAGCACACA.

#### RaPID procedure for the precipitation of RNP complexes

The pulldown of MS2 aptamer-tagged mRNAs and detection for bound RNAs and proteins was performed using the RaPID procedure, essentially as described (Slobodin & Gerst, 2010, 2011b). Yeast strains bearing endogenously expressed genes tagged with the MS2 aptamer created based on the protocol detailed in (Haim-Vilmovsky & Gerst, 2009) were grown in a volume of 400 ml to mid-log phase at 26°C with constant shaking to an O.D.600=∼0.8. Cells were transferred to preheated medium and then subjected to heat shock at 50℃ for 10 min. Cells were centrifuged in a Sorvall SLA3000 rotor at 1100 xg for 5 min, resuspended in 200 ml of complete synthetic medium lacking methionine to induce the expression of the MS2-CP-GFP-SBP protein, and grown for an additional 45 min. The cells were collected by centrifugation as described above, washed with PBS buffer (lacking Ca^++^ and Mg^++^), and transferred into a 50 ml tube and pelleted as above. Proteins were crosslinked by the addition of 8 ml PBS containing 0.01% formaldehyde and incubated at room temperature for 10 min with slow shaking. The crosslinking reaction was terminated by adding 1 M glycine buffer, pH = 8.0, to a final concentration of 0.125 M with additional shaking for 2 min. The cells were then washed once with ice-cold PBS buffer, and the pellet was flash frozen in liquid nitrogen and stored at −80°C.

For cell lysis and RNA pulldown, cell pellets were thawed upon the addition of ice-cold lysis buffer (20 mM Tris-HCl at pH 7.5, 150 mM NaCl, 1.8 mM MgCl2, and 0.5% NP40 supplemented with aprotinin [10 mg/ml], phenylmethylsulfonyl fluoride (PMSF) [1 mM], pepstatin A [10 mg/ml], leupeptin [10 mg/ml], 1 mM dithiothreitol (DTT), and 80 U/ml RNAsin [Promega]) at 1 ml per 100 O.D.600 U, and 0.5 ml aliquots were then transferred to separate microcentrifuge tubes containing an equal volume of 0.5 mm glass beads, and vortexed in an Vortex Genie Cell Disruptor (Scientific Instruments, New York) shaker at maximum speed for 10 min at 4°C. Glass beads and unbroken cells were sedimented at 4°C by centrifugation at 1700 x *g* for 1 min, and the supernatant was removed to new microcentrifuge tubes and centrifuged at 11,000 x *g* at 4°C for 10 min. The total cell lysate (TCL) was then removed to a fresh tube and protein concentration was determined using the microBCA protein determination kit (Pierce). Protein (10 mg of total cell lysate) was taken per pulldown reaction. To block endogenous biotinylated moieties, protein aliquots were incubated with 10 μg of free avidin (Sigma) per 1 mg of input protein for 1 hr at 4°C with constant rotation. In parallel, streptavidin-conjugated beads (Streptavidin-Sepharose high performance, GE Healthcare) were aliquoted to microcentrifuge tubes according to 5 µl of slurry per 1 mg of protein (not >30 μl overall), washed twice with 1 ml of PBS, once with 1 ml of lysis buffer, and blocked with a 1:1 mixture of 1 ml of lysis buffer containing yeast tRNA (Sigma; 0.1 mg/100 ml of beads) and 1 ml of 4% BSA in PBS at 4°C for 1 h with constant rotation. Following blocking, beads were washed twice in 1 ml of lysis buffer. Pulldown was performed by adding the indicated amount of avidin-blocked TCL to the beads, followed by incubation at 4°C for 2–15 hrs with constant rotation. Yeast tRNA was added to the pulldown reaction (0.1 mg/tube) to reduce nonspecific interactions. We used standard 1.7 ml microcentrifuge tubes when working with small volumes of TCL or 15 ml sterile polypropylene centrifuge tubes with larger volumes. Following pulldown, the beads were centrifuged at 660 x *g* at 4°C for 2 min, the supernatant then removed, and the beads washed three times with lysis buffer (e.g., 1 ml volume washes for small tubes, 2 ml for large tubes), twice with wash buffer (20 mM Tris, pH 7.5, 300 mM NaCl, and 0.5% Triton X-100), all performed at 4°C (each step lasting 10 min with rotation). The beads were then equilibrated by a final wash in 1–2 ml of cold PBS, pelleted by centrifugation as above, and excess buffer aspirated. For elution of the crosslinked RNP complexes from the beads, 150 ml of PBS containing 6 mM free biotin (Sigma) was added to the beads, followed by 1 h of incubation at 4°C with rotation. After centrifugation at 660 x *g* for 2 min, the eluate was transferred into a fresh microcentrifuge tube, re-centrifuged, and transferred into another tube to assure that no beads were carried over. To reverse the crosslink, the eluate was incubated at 70°C for 1–2 h with an equal volume of 2X crosslink reversal buffer (100 mM Tris, pH7.0, 10 mM EDTA, 20 mM DTT, and 2% sodium dodecyl sulfate [SDS]) for RNA analysis or with an appropriate volume of 5X protein sample buffer (5X: 0.4 M Tris at pH 6.8, 50% glycerol, 10% SDS, 0.5 M DTT, and 0.25% bromophenol blue) for protein analysis using SDS-PAGE. For RaPID pulldowns and RT-PCR analysis, three biological replicas were performed each and the average ± standard deviation is given. Primers used for qPCR reactions are listed in Supplementary File 1.

## Acknowledgements

We kindly thank Robert Singer (Albert Einstein College of Medicine, New York, USA) for providing the HSPA8 smFISH probes, Maria Vera Ugalde (McGill University, Canada) for the human HSF1-GFP lentivirus vector, Thomas Czerny (Department of Applied Life Sciences, University of Applied Sciences, Vienna, Austria) for the pGLuc and 6xHSE-Rluc plasmids, and Ruth Scherz-Shouval (Weizmann Institute of Science, Israel) for the HSF^-/-^ MEFs. Thanks also to Miriam Waghalter and Anand Govindan Ravi for EF1α-HSF1-GFP plasmid construction and lentiviral packaging, and Gal Haimovich for lentiviral transduction and FACS sorting. This work was supported by grants to J.E.G from the University of Michigan-Israel Partnership for Research and Education Foundation and from the Minerva German-Israel Foundation.

## Supplementary Figures

**Supplementary Figure 1.**
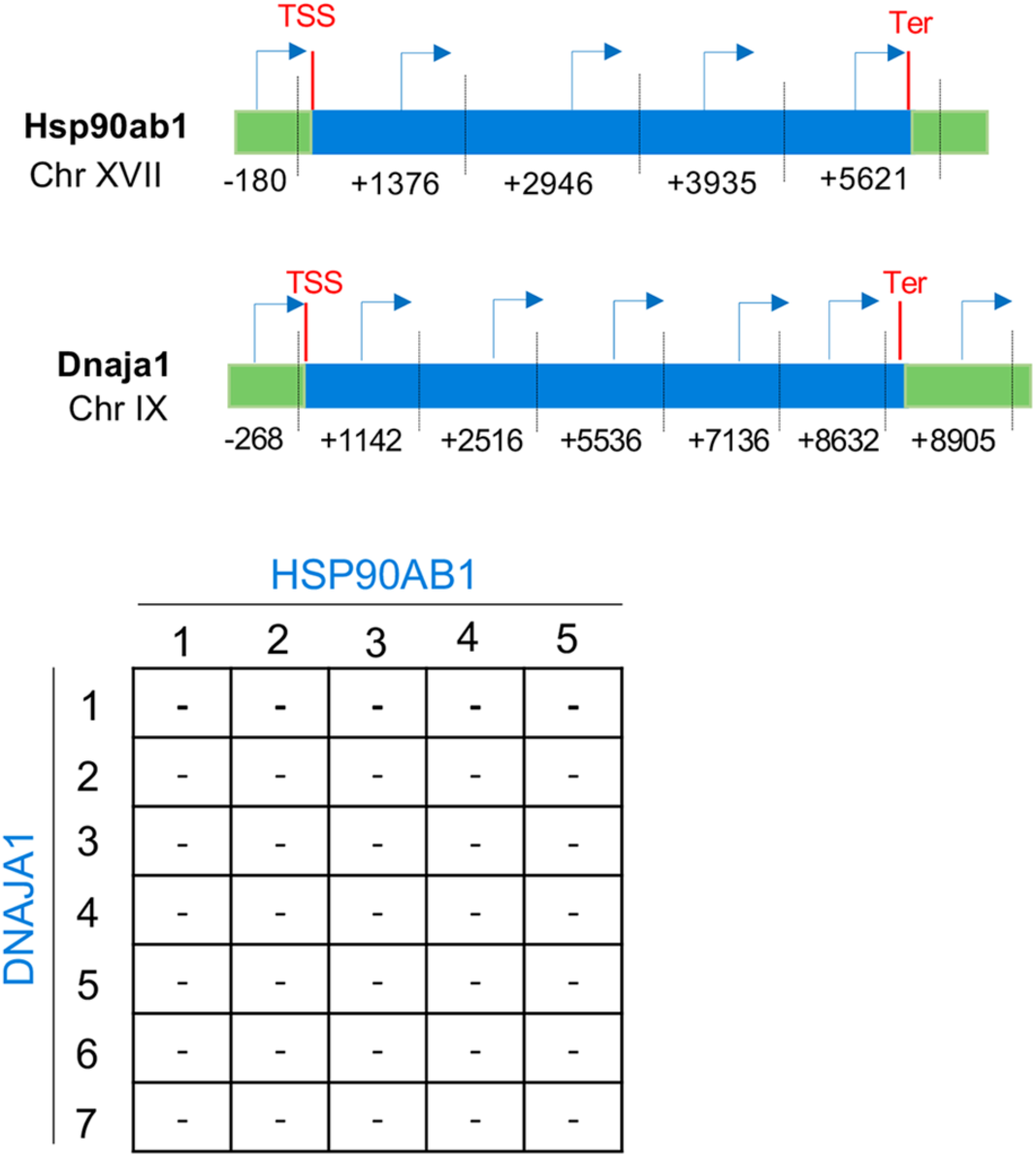
No interaction is observed between the HSP90AB1 and DNAJA1 genes upon heat shock in MEFs, as determined by the Chromosome conformation capture (3C) technique. Top panel: Graphic illustration of the HSP90AB1 and DNAJA1 genes, and the oligonucleotides used for their amplification from 3C DNA samples. Coordinates correspond to TaqI sites (shown as vertical dashed bars); site numbering is relative to ATG (+1). Forward primers used for 3C analysis were sense-strand identical (arrows) and positioned proximal to TaqI sites as indicated. Primers are numbered to distinguish the pairs used in the PCR reactions shown in matrix. 5’UTRs, ORFs, and 3’UTRs are color-coded, as indicated. Transcription start sites (TSS) and termination sites (Ter) are indicated. Bottom panel: A matrix summarizing the intergenic associations of represented genes as determined by 3C-PCR. Primer pairs corresponding to the different genes listed in were used in PCR reactions. ‘–’ indicates no amplification. *GAPDH* primers were used as a control for the PCR reaction (not shown).\

**Supplementary Figure 2.**
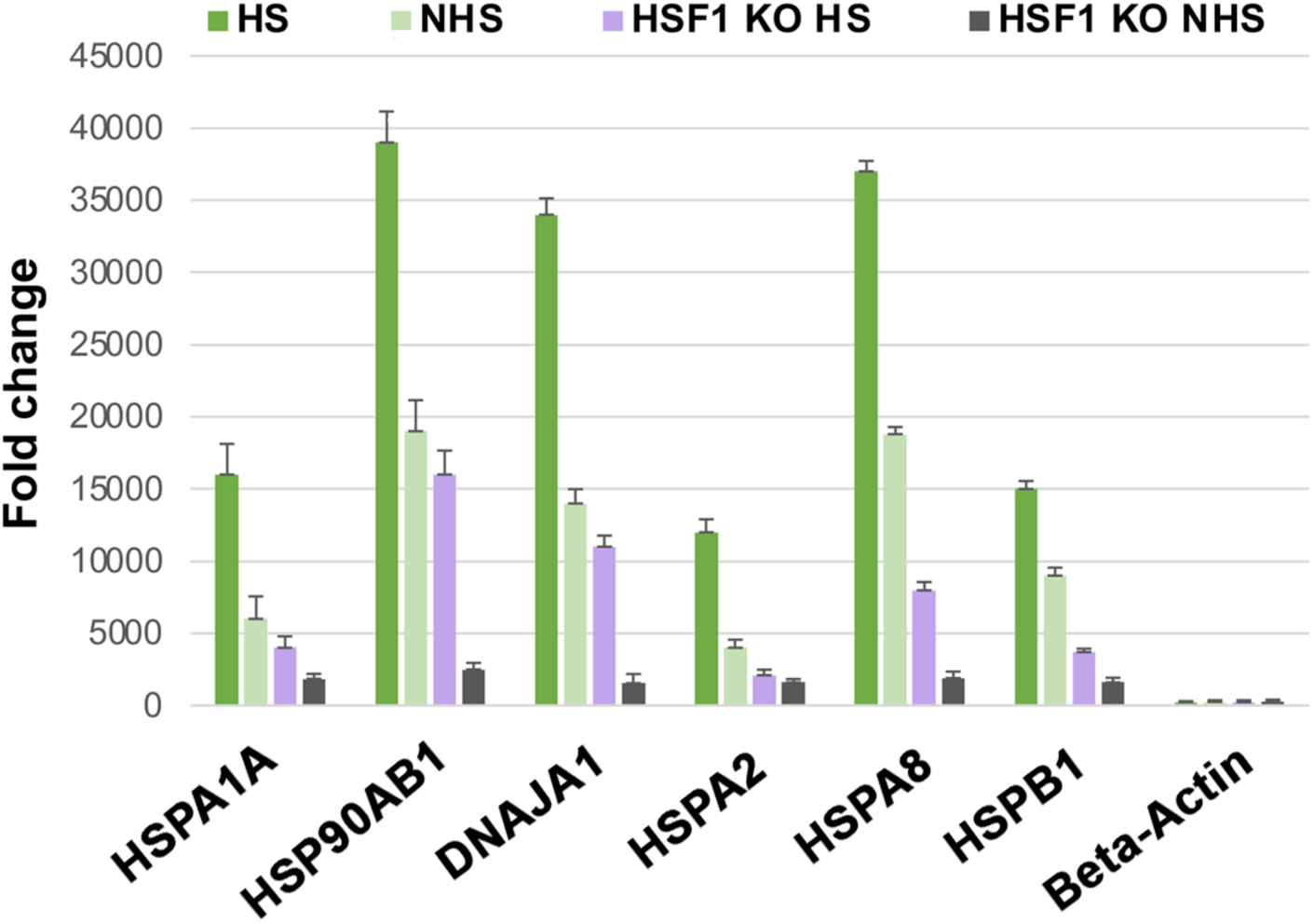
Expression levels of the different HSP genes in heat-shocked and non-heat shock exposed cells. Both WT MEFs and HSF1^-/-^ KO MEFs were exposed to heat shock for 1 hr at 42℃ or maintained at 37℃, before RNA extraction and quantification of the mRNA expression levels by qRT-PCR using specific oligos for the genes listed. Three independent replicas were performed.

**Supplementary Figure 3.**
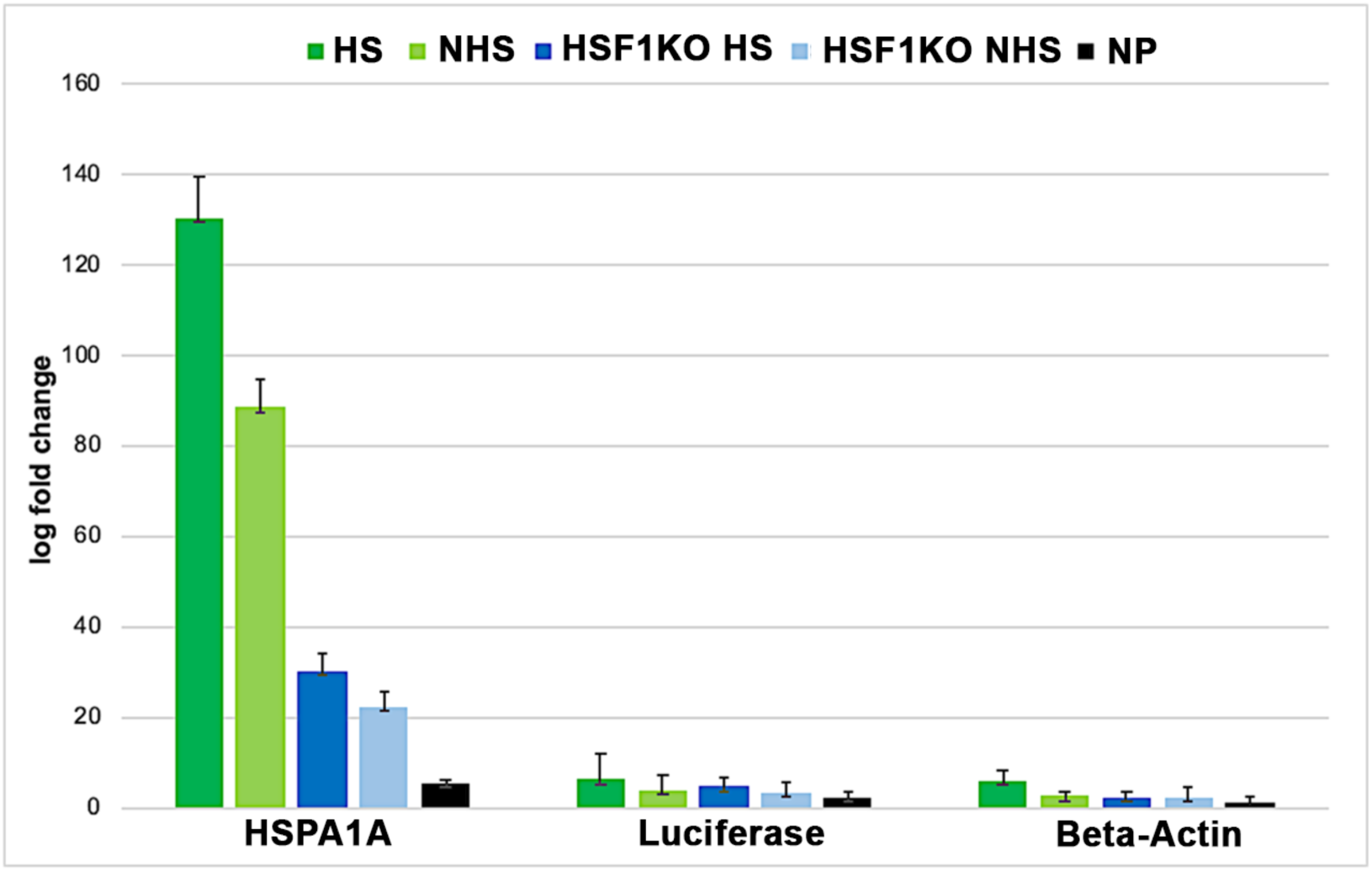
HSF1 and the HSP promoter containing a HSRE drive HSP RNA multiplexing. pGLuc plasmid was transiently expressed in WT and HSF1^-/-^ KO MEF cells and either subjected to heat shock at 42℃ for 1 hr or maintained at 37 ℃ prior to fixation and pulldown procedure. RT-PCR was performed using specific oligos against the listed genes and quantified after normalization for expression Three biological replicates were performed.

## Notes

### Competing Interest Statement

The authors have declared no competing interest.

